# Triple Combination Nitazoxanide, Ribavirin, and Hydroxychloroquine results in the multiplicative reduction of *in vitro* SARS-CoV-2 viral replication

**DOI:** 10.1101/2020.11.25.399055

**Authors:** Elena Lian, Carley McAlister, Gabriela Ramirez, David N. Chernoff, Gregory Went, Justin Hoopes, Rushika Perera

**Affiliations:** Center for Vector-borne Infectious Diseases, Department of Microbiology, Immunology, and Pathology, Colorado State University, Fort Collins, CO, USA; Synavir Corporation, San Francisco, CA, USA

**Keywords:** severe acute respiratory syndrome coronavirus-2, SARS-CoV-2, nitazoxanide, ribavirin, hydroxychloroquine

## Abstract

**Background:** An immediate unmet medical need exists to test and develop existing approved drugs against SARS-COV-2. Despite many efforts, very little progress has been made regarding finding low-cost oral medicines that can be made widely available worldwide to address the global pandemic.

**Methods:** We sought to examine if a triple combination of nitazoxanide (using its active metabolite tizoxanide), ribavirin, and hydroxychloroquine would lead to a multiplicative effects on viral replication of SARS-COV-2 resulting in a significant reduction of virus yield using VERO E6 cells as a model of viral replication.

**Results:** Virus yield measured in PFU/ml was ~ 2 logs lower with triple combination versus either drug alone, resulting in the prolongation of time to peak cytopathic effects (CPE). The time to produce 50% CPE increased from 2.8 days for viral controls versus 5.3 days for triple combination therapy. Finally, for each 1-log reduction in virus yield 24 hours post-infection, there was an additional 0.7-day delay in onset of CPE.

**Conclusions:** A triple combination of tizoxanide, ribavirin, and hydroxychloroquine produced a reduction in SARS-COV-2 viral replication in Vero E6 cells, warranting exploration in additional cell lines as well as human clinical trials.

## Introduction

From its origins in late 2019 in Wuhan, China, infection with severe acute respiratory syndrome coronavirus-2 (SARS-CoV-2) has rapidly become a global pandemic [1]. The clinical manifestations of coronavirus disease-2019 (COVID-19) range from asymptomatic carriage and mild upper respiratory symptoms to severe pneumonia, respiratory failure, sepsis, and death [2]. Viral transmission occurs predominantly via inhalation or by contact with infectious droplets [3]. In addition to spreading from symptomatic persons, pre-symptomatic and asymptomatic hosts also can transmit SARS-CoV-2, complicating efforts to contain its spread [4].

Efforts to mitigate the spread of the virus have focused on buying time using social distancing and masks to reduce the overall impact of a surge of admissions to hospitals and medical resource utilization [5]. Longer-term efforts are focused on prevention by raising herd immunity through vaccination. As of November 2020, there are over 48 vaccines in clinical development, with two of the most advanced vaccines showing high-level protection (>90%) in completed Phase 3 trials [6–8].

Despite these significant advances in vaccine-induced protection, it is impossible to predict how durable the immune response will be, with recent data suggesting durability may be limited to approximately 3 to 6 months [9]. Moreover, extremely complex logistical issues combined with biases against vaccines will hamper the distribution and achievement of meaningful coverage of the global population in the near future [10]. Hence the need for antiviral treatments for COVID-19, especially oral therapies, that can be made available to the general population in order to reduce viral load, viral spread, clinical symptoms, hospitalizations, and mortality.

Experience treating patients admitted to the hospital with influenza with a high viral load at presentation suggests that a combination of multiple antiviral drugs is more effective than single-drug treatments [11]. For influenza, it was previously discovered that selecting drugs that block *sequential steps* in viral replication could lead to synergy or a *multiplicative increase* in antiviral potency. The triple combination therapy of amantadine, ribavirin, and oseltamivir was 20-fold synergistic in both *in vitro* and *in vivo* models [12]. Results from a global Phase 2 clinical trial showed that this three-antiviral drug combination significantly reduced viral load compared to monotherapy (i.e., oseltamivir alone) [13]. A similar finding was observed for SARS and SARS-CoV-2 infections, where a double combination of interferon β-1b combined with ribavirin was shown to have a synergistic benefit *in vitro* [14, 15], as well as a significant reduction in both viral load and/or symptoms in clinical trials [16, 17]. Notably, while neither drug has a measurable benefit in clinical studies alone, taken together, they can suppress both viral load and symptoms.

For this study, we selected three FDA-approved drugs (nitazoxanide, ribavirin, and hydroxychloroquine) with different intracellular targets, two of which (ribavirin and hydroxychloroquine) have been previously evaluated in the clinic for anti-coronavirus activity but failed to show any clinical benefits when used as a monotherapy. Our goal was to determine if the combination of existing drugs that putatively block sequential steps in the replication process would lead to a multiplicative reduction in virus load *in vitro* compared to either drug alone.

## Materials & Methods

### Reagents

Tizoxanide (TIZ; CAS 173903-47-4; Sigma-Aldrich CAT# ADVH0430B80D), an active metabolite of nitazoxanide (NTZ), was prepared in DMSO, ribavirin (RBV; CAS 36791-04-5; Sigma-Aldrich CAT# R9644), and hydroxychloroquine sulfate (HCQ; CAS 747-36-4; Sigma-Aldrich CAT# PHR1782-1G) were prepared in ultrapure water and sterile filtered before use.

### Cells and Virus

SARS-CoV-2/WA/20/01 (GenBank MT020880) was acquired from the CDC. African green monkey kidney (Vero, subtype E6) cells were obtained from ATCC (CRL-1586). Cells were cultured, split, and seeded in high glucose Dulbecco’s Modified Eagle’s Medium (DMEM) (ThermoFisher cat. no. 12500062) supplemented with 2mM L-glutamine (Hyclone cat. no. H30034.01), non-essential amino acids (Hyclone cat. no. SH30238.01), 10% heat-inactivated Fetal Bovine Serum (FBS) (Atlas Biologicals cat. no. EF-0500-A), and 25 mM HEPES. Cells were seeded overnight in 6-well plates at 1.2 × 10^6^ cells/well and cultured at 37° C in a humidified incubator with 5% CO2. Cell culture and drug prep were performed under biosafety level 2 conditions while all work involving infected cells took place under biosafety level 3 conditions.

### Study Design

Vero E6 cells were washed with DMEM and treated with either 0.32 mcg/mL TIZ, 0.32 mcg/mL HCQ, or 100 μg/mL RBV as single agents as well as in double and triple combination before infection. Each condition was run in triplicate. Drug concentrations were selected based on preliminary experiments and represent concentrations that approximate human physiologic concentrations, with sub-optimal activity as single agents. Cells were washed once with DMEM and then treated for 4 hours with media containing 2% FBS and each drug alone or in combination for 4 hours. Drug pretreatment was extended because RBV uptake takes several hours in Vero E6 cells, after which it is slowly converted by adenosine kinase to the active metabolite RBV-MP [18,19]. Following pre-treatment, each test well was infected with 10 plaque-forming units (PFUs) of SARS-CoV-2 (multiplicity of infection (MOI) of <0.00001). The very low MOI allowed for the maximum number of rounds of reinfection. Cell culture supernatant samples were collected every 24 hours post-infection (PI) for 6 days and visually scored for cytopathic effects (CPE) at the time of sample collection.

The impact of drugs on virus yield in PFU/mL was determined for each collected sample by plaque assay. Briefly, 12-well plates were seeded with Vero E6 at 6 x 10^5^ cells/well and allowed to adhere overnight. The next day, cells were washed with DMEM and infected with 200 mcL of sample diluted from 10^-1^ up to 10^-6^ for 1 hour. Following absorption, plates were covered with DMEM supplemented with 5% FBS + 1% Agarose. 48 hours after initial infection, 1 mL of 0.33% Neutral Red diluted 1:24 in 1X PBS was added to the top of the agarose media and allowed to stain for an additional 24 hours. The stain was then removed, and the number of plaques were counted in each well. The first dilution that resulted in countable plaques was used to calculate the virus yield in PFU/mL.

### Statistical Analysis

Mean and standard deviation for virus yield was calculated and plotted using Graphpad Prism Ver.8.3.1. The sigmoidal curve was fit to a Hill equation, with Hill coefficient set to 1.0,[20] CPE max set to 100%, allowing for the estimation of the time required to see 50% CPE in days post-infection for each curve. Values from each replicate curve were exported to EXCEL to calculate standard deviations.

## Results

Viral yield changes were consistent across all three drugs tested and showed an increasing effect upon adding one drug to another (Figure 1 a-c). Viral yield was reduced the most for the triple combination of TIZ, RBV, and HCQ, followed by all two-drug combinations then by each single compound alone. At 24 hours post-inoculation (PI), virus yield was 3.7 ± 0.1 log10 PFU/mL in the virus control versus 2.9 ± 0.1, 2.8 ± 0.6, and 1.8 ± 2 log10 PFU/mL, in 0.32 mcg/mL TIZ, 0.32 mcg/mL HCQ, 100 mcg/mL RBV groups, respectively (Figure 2a). The combination of TIZ and HCQ, HCQ and RBV, TIZ, and RBV produced 2.3 ± 0.2, 1.9 ± 0.4, and 1.0 ± 0.9 log10 PFU/mL, respectively. The combination of all 3 drugs produced 0.8 ± 0.7 PFU/mL. Virus yield for each condition at 48 hours continued in a similar fashion with 6.0 ± 0.1 log10 PFU/mL in the virus control versus 5.3 ± 0.3, 5.2 ± 0.2, and 3.8 ± 0 log10 PFU/mL in the 0.32 mcg/mL TIZ, 0.32 mcg/mL HCQ, 100 mcg/mL RBV, respectively (Figure 2b). The combination of TIZ and HCQ, HCQ and RBV, TIZ, and RBV produced 4.5 ± 0.7, 3.7 ± 0.2, and 3.3 ± 0.1 log10 PFU/mL, respectively. The combination of all 3 drugs produced 2.4 ± 0.8 PFU/mL.

**Figure 1.**
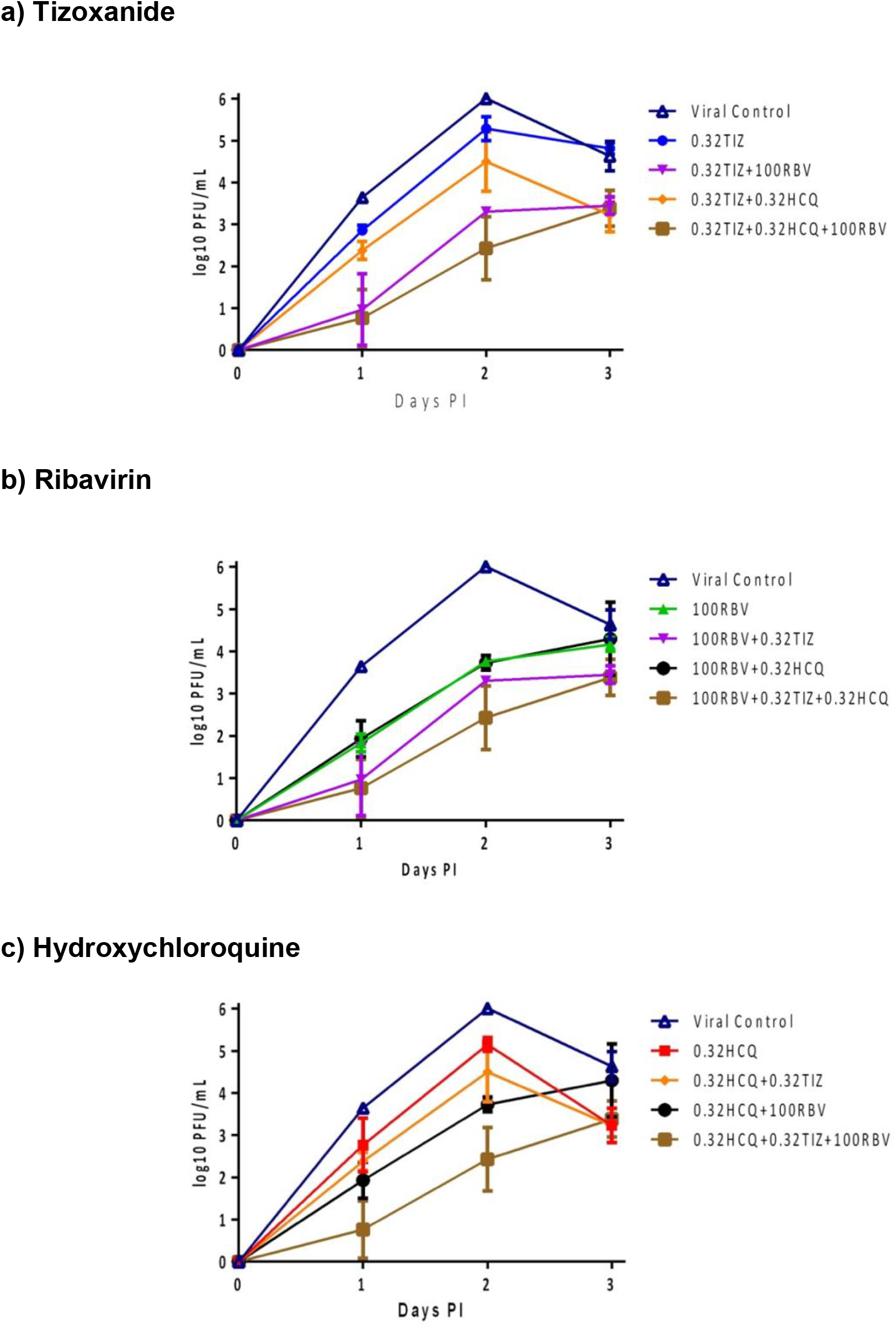
Virus yield change measured in PFU/mL by day, for each condition through day 3; a) TIZ, alone and in combination; b) RBV, alone and in combination, and c) HCQ, alone and in combination.

**Figure 2.**
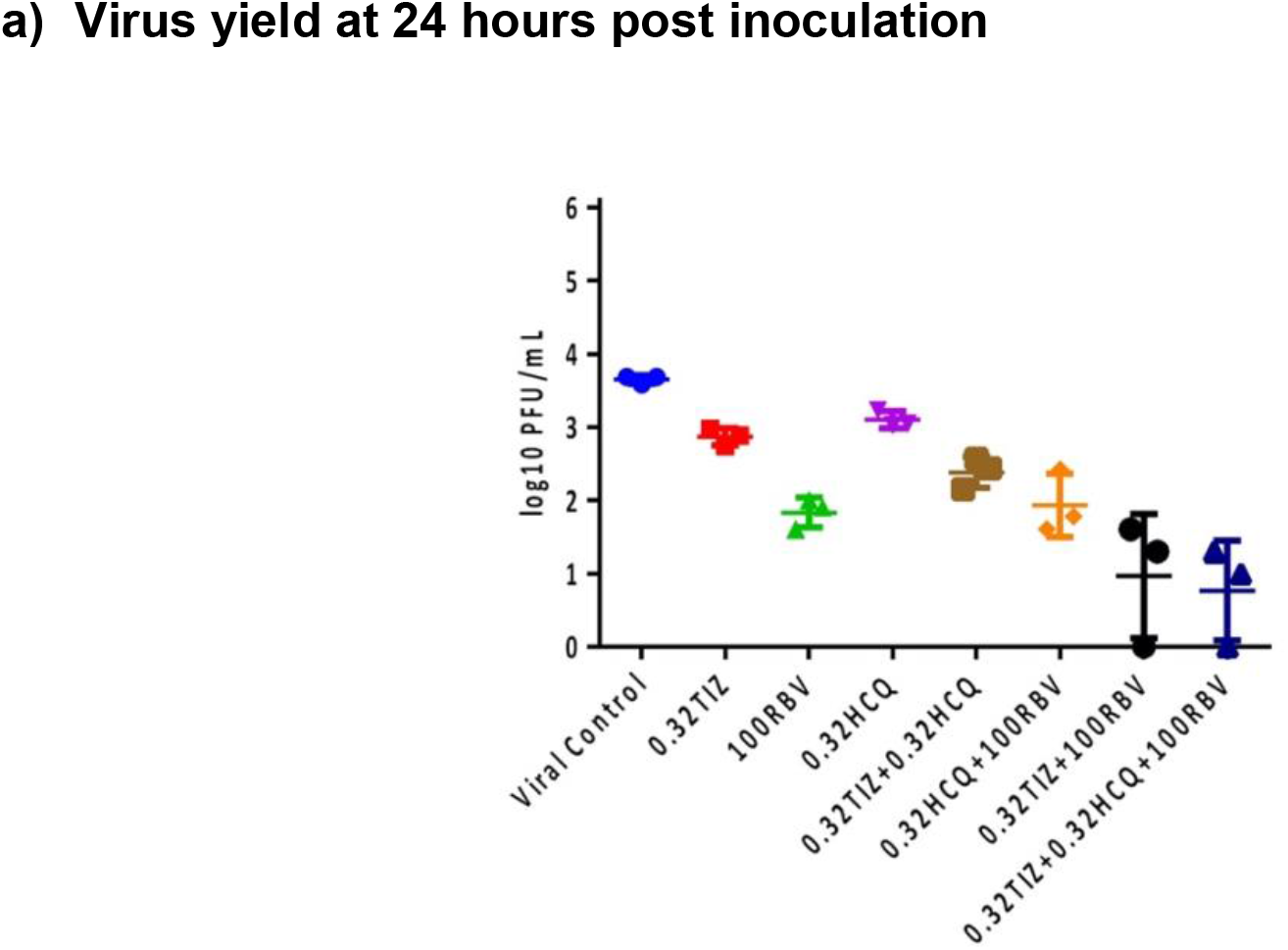
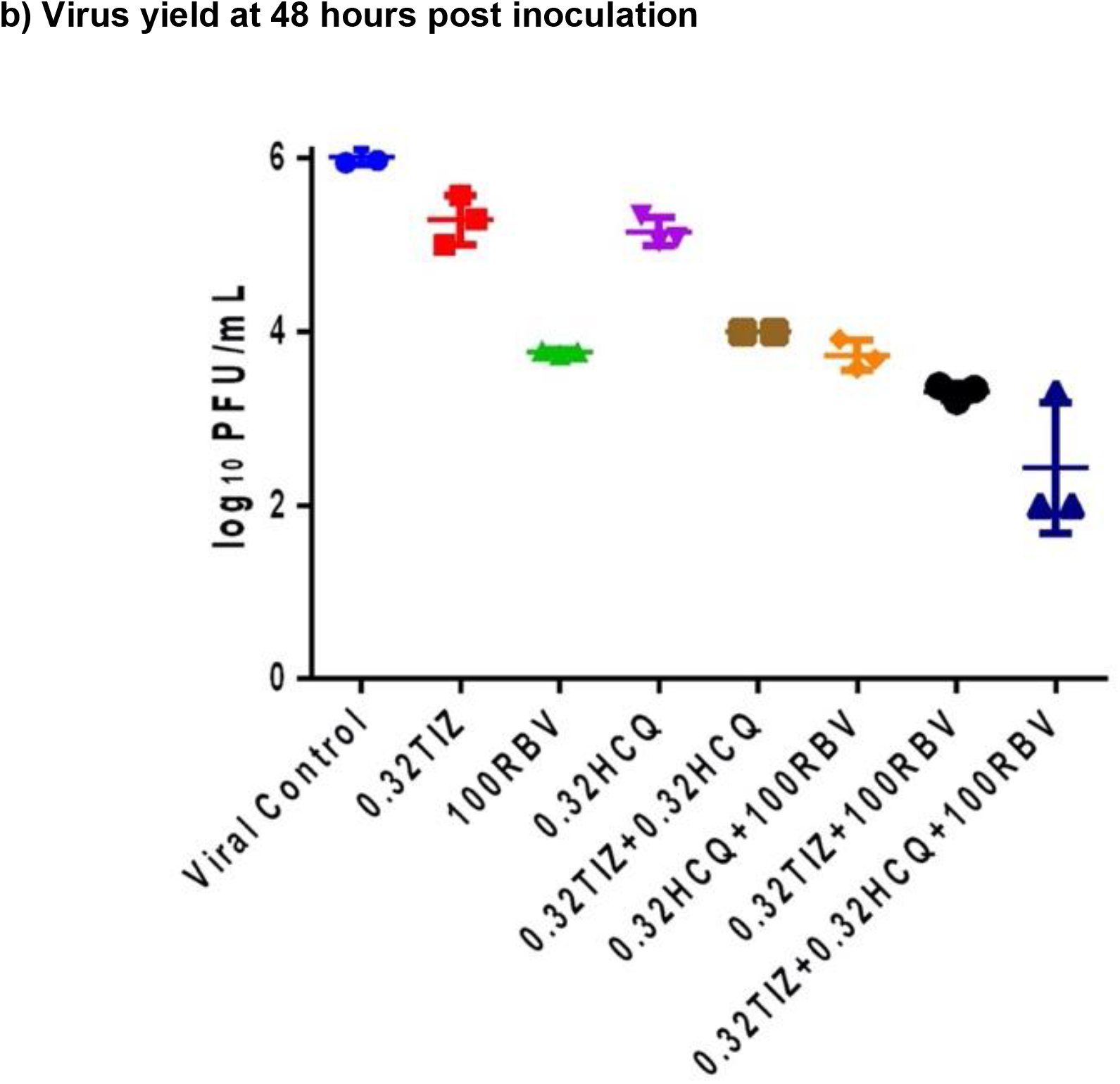
**a)** Virus yield for each condition at 24 hours post inoculation. Virus control (VC, Blue Circle) had 3.7 ± 0.1 log10 PFU/mL. 0.32 mcg/mL TIZ (Red Square), 100 mcg/mL RBV (Green Triangle), 0.32 mcg/mL HCQ (Purple Triangle) produced 2.9 ± 0.1, 1.8 ± 2, and 2.8 ± 0.6 log10 PFU/mL, respectively. The combination of TIZ and HCQ (Brown Square), HCQ and RBV (Orange Diamond), TIZ and RBV (Black Circle) produced 2.3 ± 0.2, 1.9 ± 0.4, and 1.0 ± 0.9 log10 PFU/mL, respectively. The combination of all 3 drugs produced 0.8 ± 0.7 PFU/mL (Dark Blue Triangle). **b)** Virus yield for each condition at 48 hours post inoculation. Virus control (VC, Blue Circle) had 6.0 ± 0.1 log10 PFU/mL. 0.32 mcg/mL TIZ (Red Square), 100 mcg/mL RBV (Green Triangle), 0.32 mcg/mL HCQ (Purple Triangle) produced 5.3 ± 0.3, 3.8 ± 0, and 5.2 ± 0.2 log10 PFU/mL, respectively. The combination of TIZ and HCQ (Brown Square), HCQ and RBV (Orange Diamond), TIZ and RBV (Black Square) produced 4.5 ± 0.7, 3.7 ± 0.2, and 3.3 ± 0.1 log10 PFU/mL, respectively. The combination of all 3 drugs produced 2.4 ± 0.8 PFU/mL (Dark Blue Triangle).

The observed effects on virus yield were compared to the predicted effects, assuming each drug makes a multiplicative contribution to the overall reduction in replication, as shown in Table 1. The double combinations of TIZ and HCQ and TIZ and RBV appear to be nearly multiplicative of the individual drug effects, whereas the HCQ and RBV are much less. The triple combination of TIZ, HCQ, and RBV appears to be close to multiplicative.

**Table 1.** Observed and predicted reduction in virus yield in combinations. The calculated values for the doubles and triples are obtained using the results from the single agents.

|  | Actual Fractional Yield | Actual Fold Reduction | Calculated Fractional Yield (Multiplicative) | Calculated Fold Reduction (Multiplicative) |
| --- | --- | --- | --- | --- |
| Virus Control | 1.00 | 1.0 |  |  |
| TIZ | 0.19 | 5.3 | - | - |
| HCQ | 0.14 | 7.2 | - | - |
| RBV | 0.0057 | 170 | - | - |
| TIZ + HCQ | 0.031 | 32 | 0.02600 | 38 |
| TIZ + RBV | 0.0020 | 500 | 0.00110 | 910 |
| HCQ + RBV | 0.0052 | 190 | 0.00079 | 1,300 |
| TIZ + HCQ + RBV | 0.00026 | 3,800 | 0.00015 | 6,600 |

The reduction seen in viral yield with drugs alone and in combination versus viral controls correlated with a prolongation of time to peak CPE, where 100% CPE was observed at day 3 onward in viral controls versus <5% at day 3 to 60% at day 6 in triple combination groups (Figure 3). TIZ, RBV, and HCQ as single agents produced 50% CPE in 3.5 days, 3.9 days, and 3.5 days, respectively. TIZ and HCQ, TIZ and RBV, and RBV and HCQ produced 50% CPE in 4.1, 4.5, and 4.6 days, versus 2.8 days for viral controls and 5.3 days for triple combination therapy (Figure 4). The relationship between changes in viral yield and onset of CPE was plotted to examine the reduction in virus yield (VYR) compared to virus control for each condition versus the time required to produce 50% CPE. Each 1-log reduction in virus yield correlated to a 0.8- day delay in onset of CPE, while a 1 log reduction at 48 hours resulted in a 0.7-day delay in onset of CPE (Figure 5).

**Figure 3:**
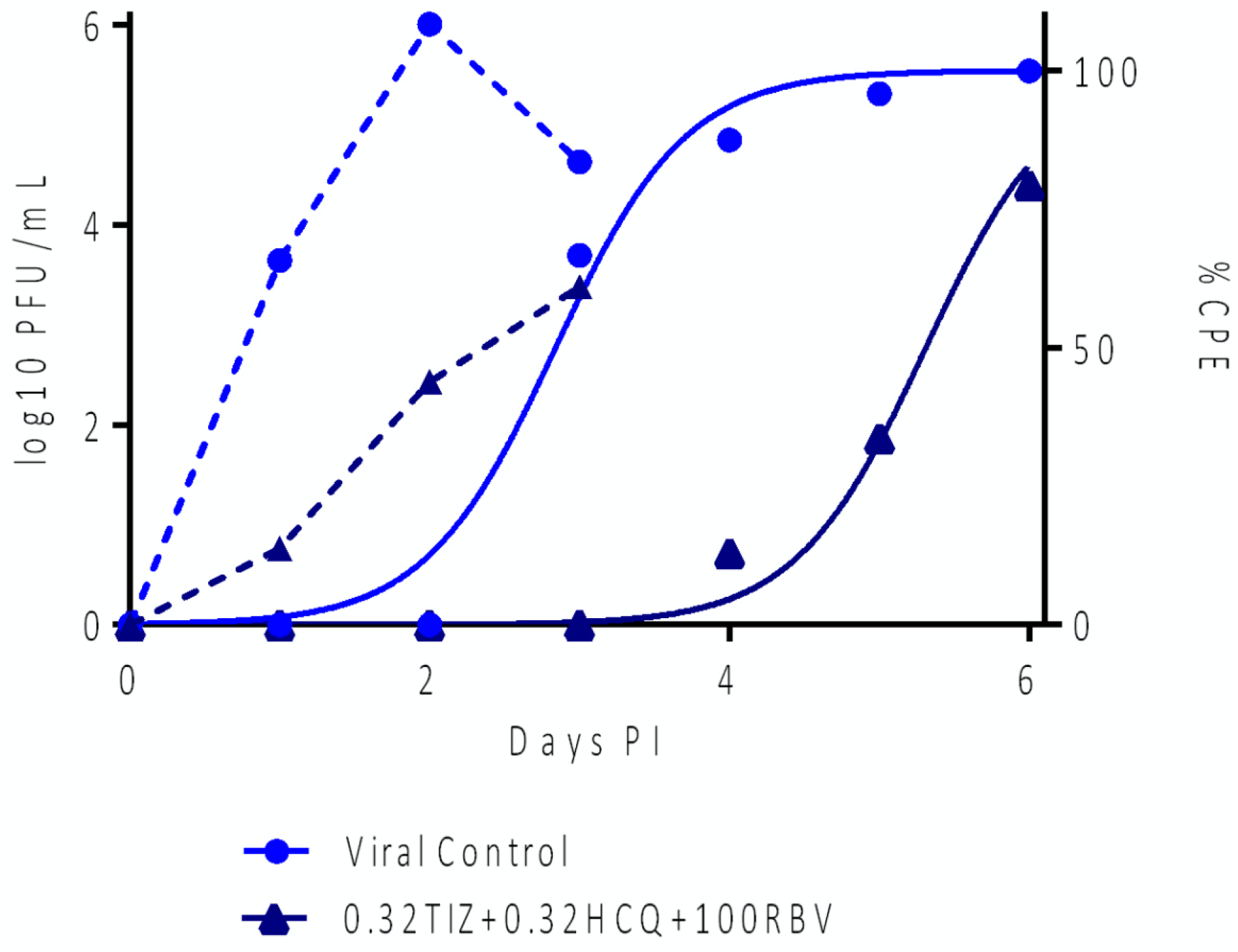
Virus yield Compared to CPE. Virus production (PFU/mL, dashed lines) and CPE (solid lines) plotted on the same graph showing the time shift between virus production and observed CPE.

**Figure 4a:**
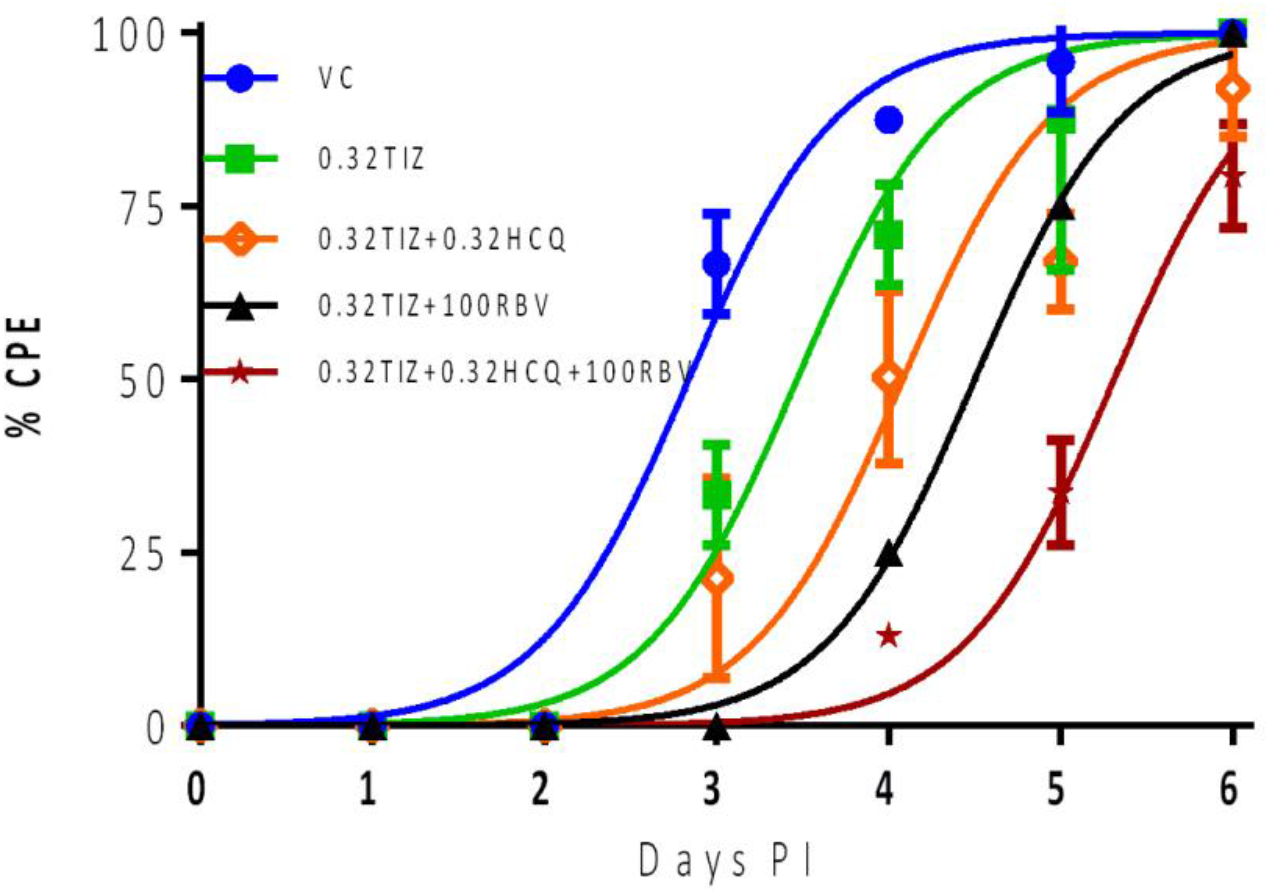
TIZ CPE as a Function of Time as a Single Agent, in Double Combination and Triple Combination. VC (Blue Circle) produced 50% CPE in 2.8 days. 0.32 mcg/mL TIZ (Green Square) produced similar CPE in 3.5 days, 0.32 mcg/mL TIZ in combination with 0.32 mcg/mL HCQ (Orange Diamond) produced similar CPE in 4.2 days, 0.32 mcg/mL TIZ in combination with 100 mcg/mL RBV (Black Triangle) produced similar CPE in 4.5 days, and 0.32 mcg/mL TIZ + 0.32 mcg/mL HCQ + 100 mcg/mL RBV (Dark Red Star) produced similar CPE in 5.3 days.

**Figure 4b:**
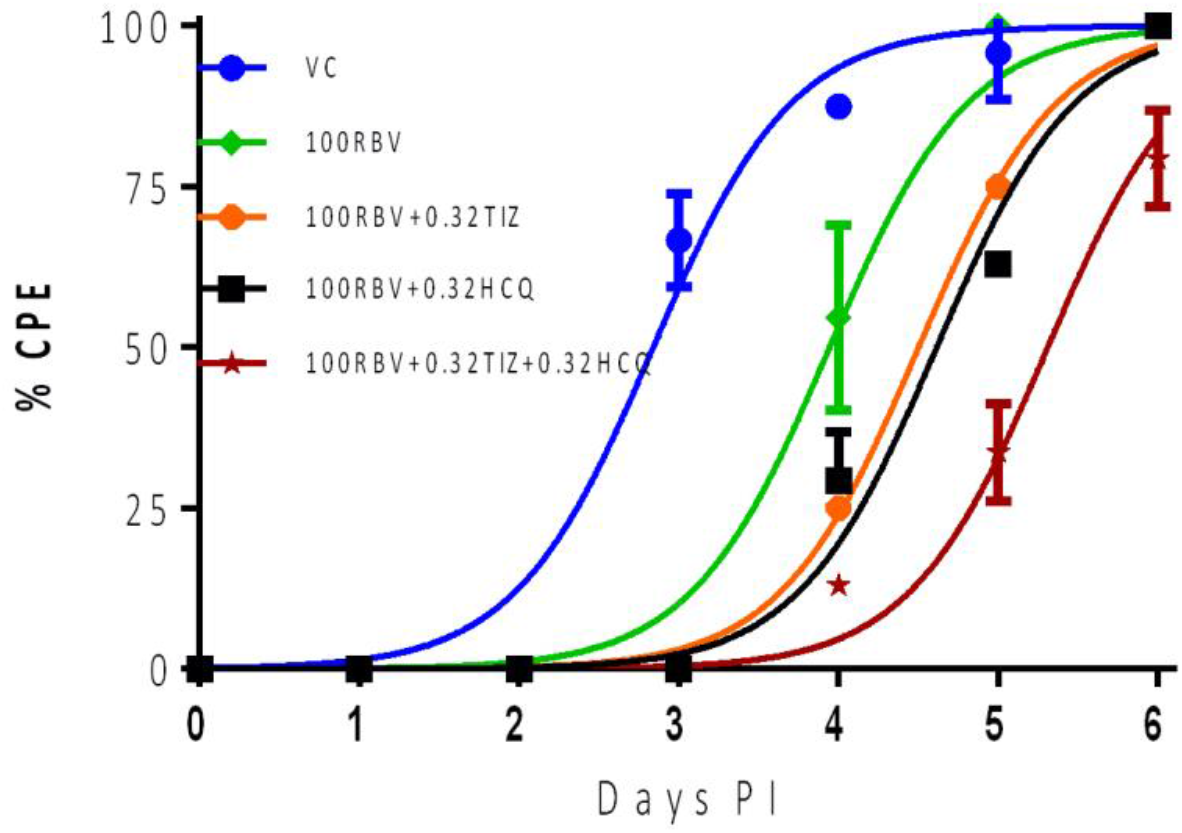
Ribavirin CPE as a Function of Time as a Single Agent, in Double Combination and Triple Combination. Virus Control (VC; Blue Circle) produced 50% CPE in 2.8 days. 100 mcg/mL RBV (Green Diamond) produced similar CPE in 3.9 days, 100 mcg/mL RBV in combination with 0.32 mcg/mL TIZ (Orange Circle) produced similar CPE in 4.5 days, 100 mcg/mL RBV in combination with 0.32 mcg/mL HCQ (Black Square) produced 50% CPE in 4.6 days while 100 mcg/mL RBV + 0.32 mcg/mL HCQ + 0.32 mcg/mL TIZ (Dark Red Star) produced similar CPE in 5.3 days

**Figure 4c:**
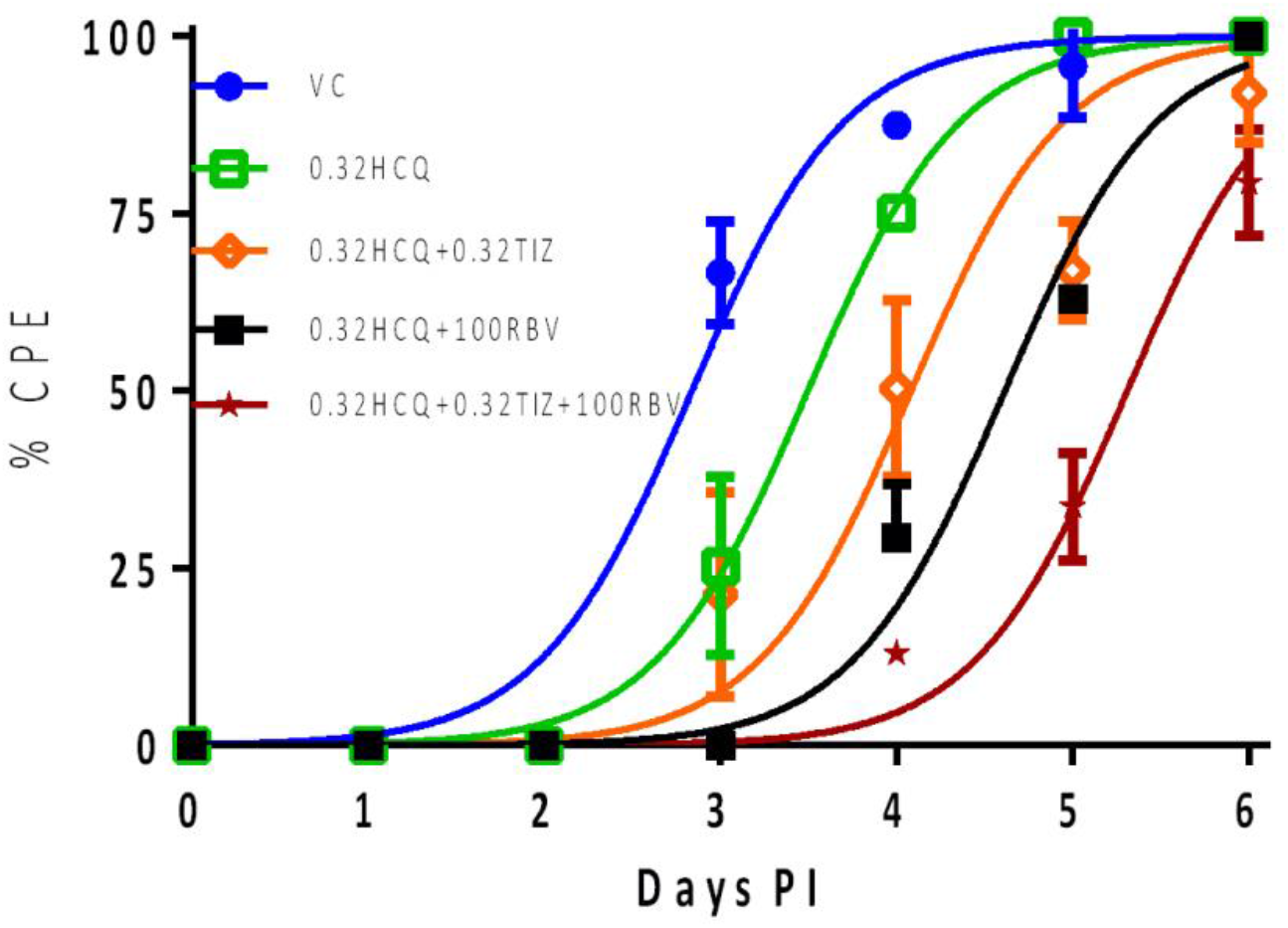
Hydroxychloroquine CPE as a Function of Time as a Single Agent, in Double Combination and Triple Combination. Virus Control (VC; Blue Circle) produced 50% CPE in 2.8 days. 0.32 mcg/mL HCQ (Green Square) produced similar CPE in 3.5 days, 0.32 HCQ in combination with 0.32 mcg/mL TIZ (Orange Diamond) produced similar CPE in 4.2 days, 0.32 mcg/mL HCQ in combination with 100 mcg/mL RBV Bblack Square) produced 50% CPE in 4.6 days while 0.32 mcg/mL HCQ + 0.32 mcg/mL TIZ + 100 mcg/mL RBV (Dark Red Star) produced similar CPE in 5.3 days.

**Figure 5:**
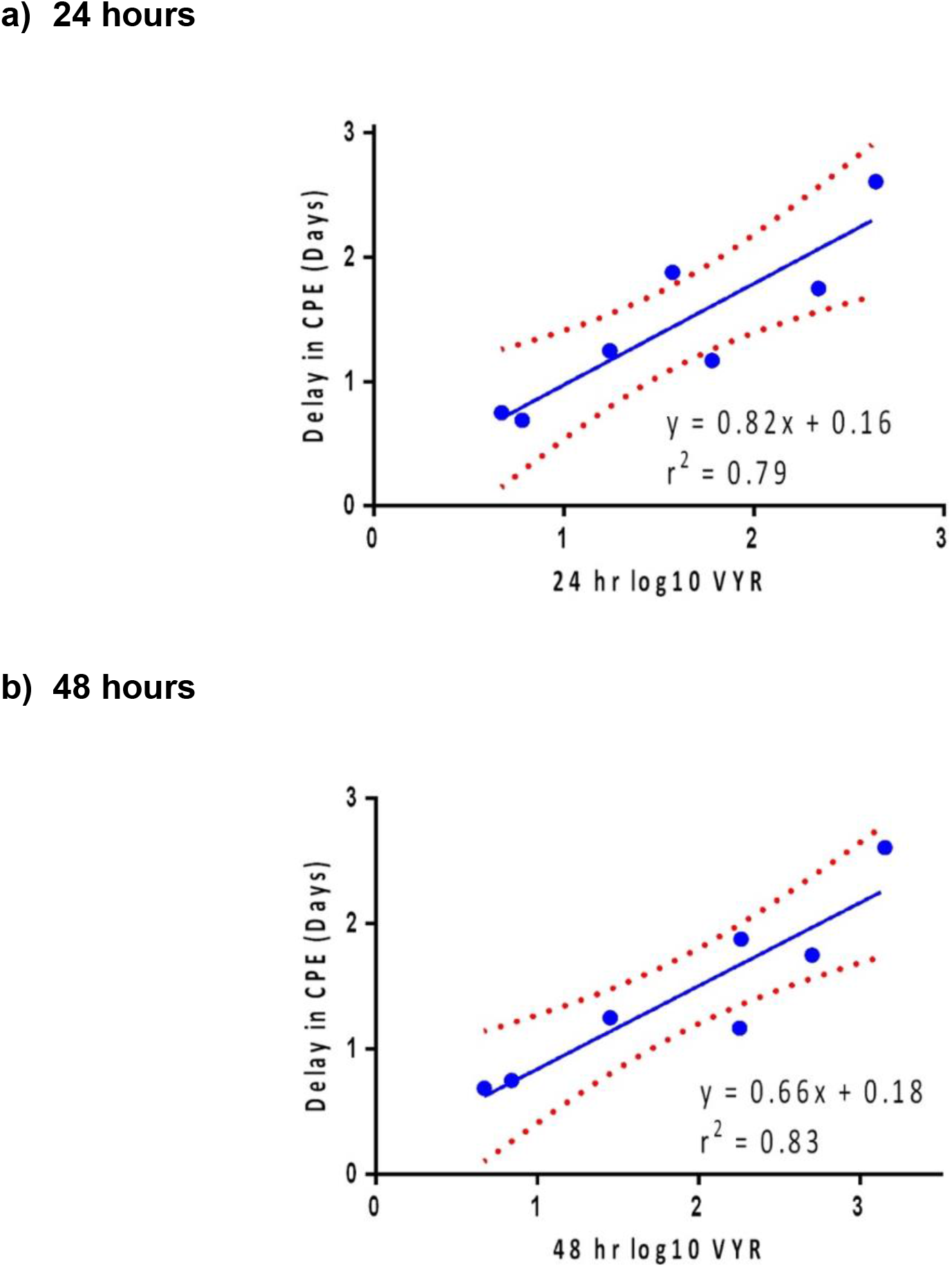
Relationship between the time required to produce 50% CPE as a function of virus yield reduction (VYR). VYR in PFU/mL at either a) 24 hours (Blue) or b) 48 hours (Blue) compared to time it takes to produce 50% CPE. Dotted red lines indicate 95% confidence interval of fit. Based on 24 hours, every 1-log reduction in CPE produces a 0.8-day delay in onset of CPE with an r^2^ of 0.79. Based on 48 hours, every 1-log reduction in CPE produces a 0.7-day delay in onset of CPE with r^2^ of 0.83.

## Discussion

Despite new data documenting that several highly protective vaccines are close to approval, there remains an immediate and sustained need for effective antiviral treatments for SARS-COV-2 infections. Several treatments are now approved for use in severe cases that have demonstrated improved outcomes in the hospital setting (remdesivir, dexamethasone). However, there are no treatments available that can be practically administered to a large outpatient population in order to reduce viral load and viral spread. Examining previously approved drugs is the fastest path to new treatments, but efforts have been hampered by lack of efficacy as monotherapies in clinical trials. We have hypothesized that combining multiple drugs targeting sequential steps in the replication process of SARS-CoV2 may help overcome the efficacy limitations observed in previous studies. To test this hypothesis, we selected three drugs, two of which (HCQ, RBV) had been previously evaluated and showed efficacy *in vitro* at physiologically achievable concentrations but failed to show effectiveness in clinical trials when used as monotherapies.

Vero E6 cells were selected for this study because SARS-COV-2 produces high titers and CPE with a fast replication rate in this cell line. A large well format with a low multiplicity of infection (MOI) was also employed to maximize the number of rounds of replication and reinfection and apply mathematical modeling to analyze the results.

Each drug was shown previously to have a dose-dependent effect on virus yield in VeroE6 cells [21,22]. In this study, drugs were administered at concentrations that corresponded to clinically achievable active metabolite concentrations. Despite being administered at sub-optimal concentrations, all three drugs reduced virus yield at days 1 and 2, with inhibitory effects ranging from 8- to 160-fold. Through day 2, the double combinations of TIZ and HCQ and TIZ and RBV further improved compared with either drug alone, each being approximately multiplicative. In contrast, the combination of RBV and HCQ did not reduce virus yield compared to RBV alone. Finally, the triple combination of TIZ, RBV, and HCQ showed a substantial reduction in virus yield compared to either drug alone or double combination, resulting in nearly a 4000-fold reduction compared to only a 160-fold and 500-fold drop in virus yield in the best monotherapy and double combination, respectively.

Compared to a simple model of combination inhibition, the results appear to exhibit a multiplicative effect. The degree of virus yield reduction is predicted by the multiplication of the decrease in individual viral yields. Further modeling work is required to differentiate between the additive and multiplicative effects of these drugs.

Not surprisingly, there was a strong association between replication rate and the onset of CPE. For every log that the virus yield was reduced at 24 hours, there was a corresponding 0.8-day delay in CPE onset. This suggests that CPE is a direct result of viral replication. These results support our primary hypothesis, namely, choosing drugs that intervene at multiple points in the replication process, potentially in series, results in a nearly multiplicative reduction in overall replication rates.

This study has several important limitations. With respect to HCQ and RBV, it has been established that Vero E6 cells are not a good proxy of antiviral activity in human airway epithelial cells. For HCQ, TMPRSS-2 expression in human airway epithelial cells may result in an alternate pathway for viral entry that bypasses the cathepsin mediated pathway used in Vero E6 cells, resulting in a much higher EC50 required to inhibit viral replication for human airway cell lines [23]. This may be offset by much higher lung concentrations for HCQ versus plasma concentration [24]. Second, RBV is very slowly taken up and subsequently phosphorylated to its monophosphate form in Vero E6 cells [18,19], resulting in high micromolar concentrations to suppress viral replication.

## Conclusions

This study demonstrated that the triple combination of tizoxanide (an active metabolite of nitazoxanide), ribavirin, and hydroxychloroquine leads to a nearly multiplicative reduction of SARS-CoV-2 *in* vitro in Vero-E6 cells. Experiments are underway to determine if the same antiviral multiplicative effect can be observed in multiple human cell lines and how this combination of drugs compares to other drugs under investigation, such as remdesivir and interferon/ribavirin. Placebo-controlled studies of the best double (NTZ and RBV) and triple combination are underway to establish the clinical benefit of combination therapy in treating COVID-19 and other human coronavirus infections (ClinicalTrials.gov identifiers: NCT04605588 and NCT4563208).

## Acknowledgements

The authors would like to thank Brian J. Geiss, Ph.D., The Office of the Vice President for Research, and the Department of Microbiology, Immunology and Pathology for assistance with establishing the antiviral testing program at Colorado State University. The authors would also like to thank Emmett Glass, Ph.D., of Red Sky Medical for providing medical writing/ editorial support during the preparation of the manuscript.

## Author statement

DC, JH, GW, and RP contributed to conceptualization and development of methodology. JH, EL, CM, GR, and RP contributed to data acquisition and analysis. DC, GW, EL, CM, GR, JH, and RP contributed to writing, reviewing, and editing.

## REFERENCES

1. Mahase E. Covid-19: Who declares pandemic because of “alarming levels” of spread, severity, and inaction. BMJ. 2020; 368:m1036.

2. Rothan HA, Byrareddy SN. The epidemiology and pathogenesis of coronavirus disease (COVID-19) outbreak. Journal of autoimmunity. 2020 Feb 26:102433.

3. Han Y, Yang H. The transmission and diagnosis of 2019 novel coronavirus infection disease(covid-19): A Chinese perspective. J Med Virol. 2020.

4. Bai Y, Yao L, Wei T, et al. Presumed asymptomatic carrier transmission of COVID-19. Jama. 2020 Apr 14;323(14):1406–7.

5. Walensky RP, Del Rio C. From mitigation to containment of the COVID-19 pandemic: putting the SARS-CoV-2 genie back in the bottle. Jama. 2020 Apr 17.

6. WHO, Draft landscape of COVID-19 candidate vaccines, 12 Nov 2020. Available at: https://www.who.int/publications/m/item/draft-landscape-of-covid-19-candidate-vaccines. Accessed 20 November 2020.

7. Pfizer and BioNTech Announce Vaccine Candidate Against COVID-19 Achieved Success in First Interim Analysis from Phase 3 Study. 2020. Available at: https://investors.biontech.de/news-releases/news-release-details/pfizer-and-biontech-announce-vaccine-candidate-against-covid-19. Accessed 20 November 2020.

8. Moderna’s COVID-19 Vaccine Candidate Meets its Primary Efficacy Endpoint in the First Interim Analysis of the Phase 3 COVE Study. 2020. Available at: https://investors.modernatx.com/news-releases/news-release-details/modernas-covid-19-vaccine-candidate-meets-its-primary-efficacy. Accessed 20 November 2020.

9. Seow J, Graham C, Merrick B, et al. Longitudinal evaluation and decline of antibody responses in SARS-CoV-2 infection. MedRxiv. Jan 1, 2020.

10. Mello MM, Silverman RD, Omer SB. Ensuring uptake of vaccines against SARS-CoV-2. New England Journal of Medicine. Jun 26, 2020.

11. Dunning J, Baillie JK, Cao B, Hayden FG. Antiviral combinations for severe influenza. The Lancet infectious diseases. 2014 Dec 1;14(12):1259–70.

12. Nguyen JT, Hoopes JD, Le MH, et al. Triple combination of amantadine, ribavirin, and oseltamivir is highly active and synergistic against drug resistant influenza virus strains in vitro. PloS one. 2010 Feb 22;5(2):e9332.

13. 3eigel JH, Bao Y, Beeler J, et al. Oseltamivir, amantadine, and ribavirin combination antiviral therapy versus oseltamivir monotherapy for the treatment of influenza: a multicentre, double-blind, randomised phase 2 trial. The Lancet Infectious Diseases. 2017 Dec 1;17(12):1255–65.

14. Chen F, Chan KH, Jiang Y, et al. In vitro susceptibility of 10 clinical isolates of SARS coronavirus to selected antiviral compounds. J Clin Virol. 2004; 31: 69–75.

15. Morgenstern B, Michaelis M, Baer PC, Doerr HW, Cinatl Jr J. Ribavirin and interferon-β synergistically inhibit SARS-associated coronavirus replication in animal and human cell lines. Biochemical and biophysical research communications. 2005 Jan 28;326(4):905–8.

16. Omrani AS, Saad MM, Baig K, et al. Ribavirin and interferon alfa-2a for severe Middle East respiratory syndrome coronavirus infection: a retrospective cohort study. The Lancet Infectious Diseases. 2014 Nov 1;14(11):1090–5.

17. Fan-Ngai HI, Kwok-Cheung L, Yuk-Keung TE, et al. Triple combination of interferon beta-1b, lopinavir-ritonavir, and ribavirin in the treatment of patients admitted to hospital with COVID-19: an open-label, randomised, phase 2 trial. Lancet. 2020; 395(10238):1695–1704.

18. Smee DF, Bray M, Huggins JW. Antiviral activity and mode of action studies of ribavirin and mycophenolic acid against orthopoxviruses in vitro. Antiviral Chemistry and Chemotherapy. 2001 Dec;12(6):327–35.

19. Shah NR, Sunderland A, Grdzelishvili VZ. Cell type mediated resistance of vesicular stomatitis virus and Sendai virus to ribavirin. PloS one. 2010 Jun 22;5(6):e11265.

20. Perelson AS, Deeks SG. Drug effectiveness explained: the mathematics of antiviral agents for HIV. Sci Transl Med. 2011 Jul 13;3(91):91ps30.

21. Dittmar M, Lee JS, Whig K, et al. Drug Repurposing Screens Reveal FDA Approved Drugs Active Against SARS-CoV-2. Available at SSRN. 2020. https://ssrn.com/abstract=3678908

22. Falzarano D, de Wit E, Martellaro C, Callison J, Munster VJ. Feldmann H. Inhibition of novel β coronavirus replication by a combination of interferon-α2b and ribavirin. Sci Rep. 2013; 3: 1686.

23. Ou T, Mou H, Zhang L, Ojha A, Choe H, Farzan M. Hydroxychloroquine-mediated inhibition of SARS-CoV-2 entry is attenuated by TMPRSS2. BioRxiv. 2020. https://doi.org/10.1101/2020.07.22.216150

24. Maisonnasse P, Guedj J, Contreras V, et al. Hydroxychloroquine use against SARS-CoV-2 infection in non-human primates. Nature 2020; 585: 584–587.

